# Biomarker-based outcome prediction in prostate adenocarcinoma depends on the *TMPRSS2-ERG* status

**DOI:** 10.1101/546200

**Authors:** Julia S. Gerke, Martin F. Orth, Yuri Tolkach, Laura Romero-Pérez, Fabienne Wehweck, Stefanie Stein, Julian Musa, Maximilian M. L. Knott, Tilman L. B. Hölting, Jing Li, Giuseppina Sannino, Aruna Marchetto, Shunya Ohmura, Florencia Cidre-Aranaz, Martina Müller-Nurasyid, Konstantin Strauch, Christian Stief, Glen Kristiansen, Thomas Kirchner, Alexander Buchner, Thomas G. P. Grünewald

## Abstract

**Background:** Prostate adenocarcinoma (PCa) with/without the *TMPRSS2-ERG* (T2E)-fusion represent distinct molecular subtypes.

**Objective:** To investigate gene-signatures associated with metastasis in T2E-positive and -negative PCa, and to identify and validate subtype-specific prognostic biomarkers.

**Design, setting and participants:** Gene expression and clinicopathological data of two discovery PCa cohorts (total *n*=783) were separately analyzed regarding the T2E-status. Selected subtype-specific biomarkers were validated in two additional cohorts (total *n*=405).

**Outcome measurements and statistical analysis:** From both discovery cohorts, we generated two gene lists ranked by their differential intratumoral expression in patients with/without metastases stratified by T2E-status, which were subjected to gene set enrichment and leading-edge analyses. The resulting top 20 gene-signatures of both gene lists associated with metastasis were analyzed for overlaps between T2E-positive and -negative cases. Genes shared by several functional gene-signatures were tested for their association with event-free survival using the Kaplan-Meier method in a validation cohort. Immunohistochemistry was performed in another validation cohort.

**Results and limitations:** Metastatic T2E-positive and -negative PCa are characterized by different gene-signatures. Five genes (*ASPN, BGN, COL1A1, RRM2* and *TYMS*) were identified whose high expression was significantly associated with worse outcome exclusively in T2E-negative PCa. This was validated in an independent cohort for all genes and additionally for RRM2 by immunohistochemistry in a separate validation cohort. No prognostic biomarkers were identified exclusively for T2E-positive tumors.

**Conclusions:** Our study demonstrates that the prognostic value of biomarkers critically depends on the molecular subtype, i.e. the T2E-status, which should be considered when screening for and applying novel prognostic biomarkers for outcome prediction in PCa.

**Patient summary:** Outcome prediction for PCa is complex. The results of this study highlight that the validity of prognostic biomarkers depends on the molecular subtype, specifically the presence/absence of T2E. The reported new subtype-specific biomarkers exemplify that biomarker-based outcome prediction in PCa should consider the T2E-status.

## INTRODUCTION

Prostate adenocarcinoma (PCa) is the second most common cancer in men worldwide, which is often detected in early stages due to regular screening [1]. Although most patients exhibit a slowly growing, indolent tumor that can be treated with active surveillance [1], 15-20% of patients develop an aggressive tumor requiring intense treatment, which is associated with significant adverse effects [2,3]. However, it remains difficult to discriminate indolent from aggressive PCa [4], wherefore 23-42% of men are ‘overtreated’ leading to unnecessary therapy-associated morbidity that may affect quality of life and life expectancy [1,5,6]. Further, overtreatment constitutes a significant socioeconomic and healthcare burden in the Western world [5]. Thus, novel strategies to discriminate aggressive from indolent disease are urgently required.

Around 50% of PCa are characterized by chromosomal rearrangements creating chimeric oncogenes through fusion of *TMPRSS2* with *ERG*, the latter belonging to the ETS family of transcription factors [7]. TMPRSS2-ERG (T2E) acts as an aberrant transcription factor with oncogenic properties [7]. Prior studies proved that T2E-positive and –negative PCa constitute molecularly distinct PCa-subtypes [8–10], which may exploit different gene-signatures or pathways to promote PCa malignancy.

A recent study highlighted the importance of certain gene-signatures for progression of PCa and suggested several genes as potential biomarkers [11]. Yet, the impact of molecular alterations such as T2E on these gene-signatures was not specifically considered.

Here, we combined transcriptome profiles and clinicopathological data of two discovery cohorts, and explored gene-signatures and their associated genes involved in metastasis depending on the T2E-status. We identified five prognostic biomarkers specifically suitable for T2E-negative PCa, which was validated in two additional cohorts. Going beyond prior studies [8,10,11], we show that the T2E-status critically determines the nature of distinct metastasis-associated gene-signatures, and strongly impacts on prognostic biomarkers.

## MATERIALS AND METHODS

### Microarray and RNA sequencing (RNA-Seq) data

Two publicly available gene expression datasets with matched clinicopathological data were downloaded from the Gene Expression Omnibus (GEO) and The Cancer Genome Atlas (TCGA). The GEO dataset (GSE46691) comprised 545 PCa cases profiled on Affymetrix GeneChip Human Exon 1.0 ST arrays [12]. Microarray signal intensities were normalized using the SCAN algorithm of SCAN.UPC [13] and the ‘pd.huex.1.0.st.v2’ annotation [14] Bioconductor packages with brainarray chip description files (CDF, huex10sthsentrez, version 21), yielding one optimized probe-set per gene (gene level summarization) [15]. The TCGA PCa dataset (TCGA-PRAD) contains preprocessed RNA-Seq level 3 data of 497 cases [10]. Based on the TNM-classification of tumors, we stratified both datasets in cases with/without metastasis (corresponding to N0M0 versus N>0 and/or M>0). As incidence and aggressiveness may be different in Africans and Afro-Americans compared to Europeans [1], we filtered – if possible – for men with European ancestry, which was carried out via principal component analysis in the TCGA-PRAD-cohort based on common SNPs identified by parallel exome sequencing. This resulted in a final TCGA-PRAD-cohort of 384 cases (**Figure 1a**).

**Figure 1.**
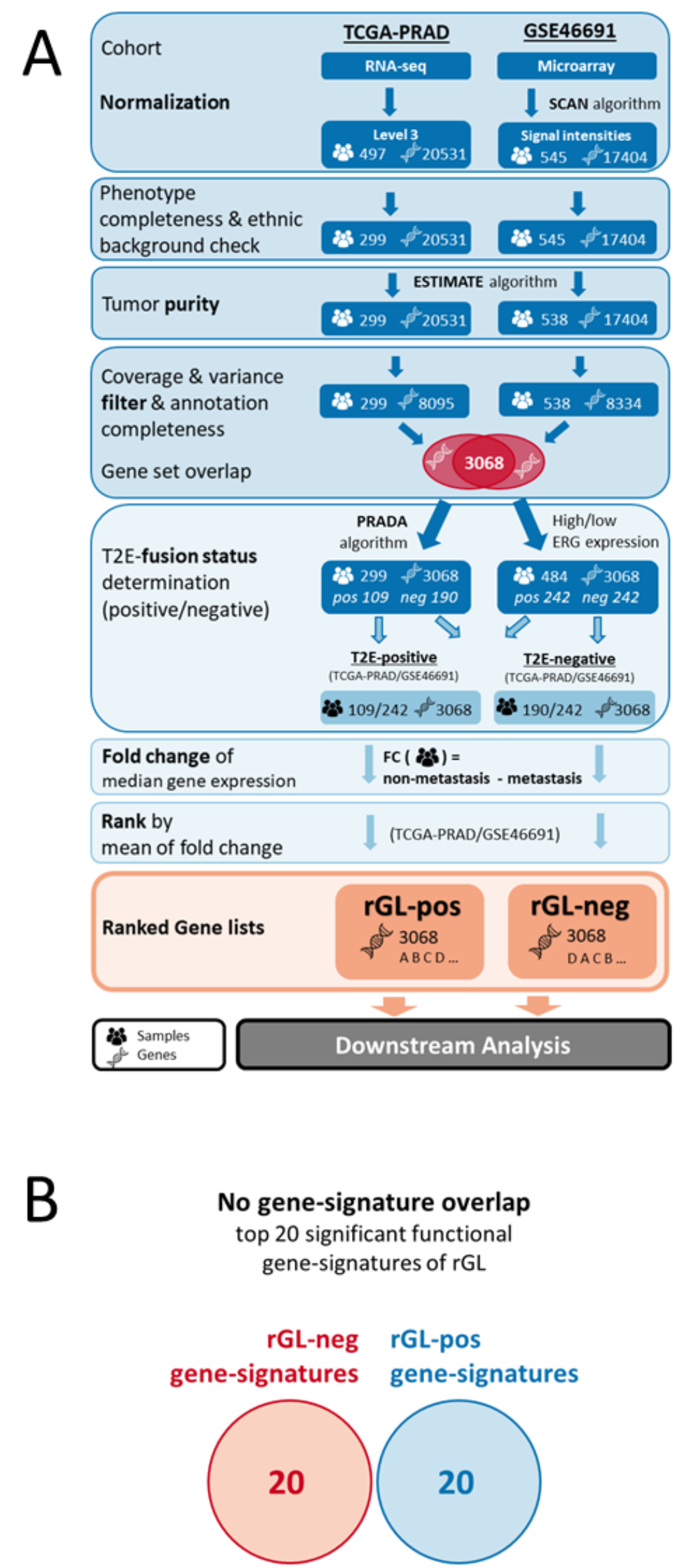
T2E-positive and -negative PCa are characterized by distinct metastasis-associated gene-signatures. **A)** Pipeline overview depicting the processing transcriptome data from the TCGA-PRAD- and, GSE46691-cohorts, and generation of differentially ranked gene lists (rGL-pos and -neg) **B)** Venn diagram showing the top 20 significant gene-signatures as identified by GSEA of rGL-pos and -neg.

### Determination of the T2E-status

In the TCGA-PRAD-cohort, the T2E-status was inferred by Torres-García *et al*. based on RNA-Seq split-reads (http://www.tumorfusions.org/) [16]. In the Affymetrix dataset (GSE46691), the T2E-status was inferred from *ERG* expression levels, which show high concordance with the T2E-status [17]. Cases were classified as T2E-positve or -negative if their individual *ERG* expression level was above/below the median *ERG* expression. To reduce the number of potentially misclassified cases, we excluded those 10% with *ERG* expression levels between the 45^th^ and 55^th^ percentile (**Supplementary Figure 1**).

### Processing of microarray and RNA-Seq data

In both cohorts, we separately determined cancer purity with the ESTIMATE algorithm [18]. Only cases with a consensus purity estimation (CPE) of >60% corresponding to TCGA standard (http://cancergenome.nih.gov/cancersselected/biospeccriteria) were kept for downstream analyses (**Supplementary Figure 2**). Next, we removed cases with <90% gene coverage and those 50% of genes with lowest variance across all samples using the genefilter Bioconductor package [19]. Moreover, transcripts or probesets from both cohorts, which could not be unambiguously annotated with official gene symbols, and genes that were represented in only one cohort were removed. The unity of both cohorts corresponded to 3,068 variably expressed genes for 299 cases from the TCGA-PRAD-cohort and 538 cases from the GSE46691-cohort. We next stratified both cohorts according to the T2E-status resulting in four sub-cohorts comprising 109 T2E-positive and 190 -negative cases for the TCGA-PRAD-cohort, and 242 T2E-positive and 242 -negative cases for the GSE46691-cohort. We then calculated in each sub-cohort separately the median fold change of each gene between samples with/without metastasis at diagnosis. Subsequently, the mean fold change from the two corresponding median fold changes was calculated for T2E-postive and -negative cases. This yielded two gene lists comprising the unity of 3,068 genes ranked by their mean fold change in T2E-positive (rGL-pos) and T2E-negative cases (rGL-neg) (**Figure 1a**).

### Gene set enrichment analysis (GSEA)

To identify significantly enriched gene-signatures (normalized enrichment score (NES) >1.6, nominal *P*<0.05 and FDR *q*<0.3) in both preranked lists (rGL-pos and rGL-neg) we employed GSEA (MSigDB v6.2; chemical and genetic perturbations; 1,000 permutations) [20].

To identify common genes across the top 20 significantly enriched gene-signatures (highest NES), we extracted those genes by leading-edge analysis that were involved in >3 gene-signatures yielding two new top gene-signature gene lists for T2E-positive and -negative cases (topGL-pos and topGL-neg) (**Figure 2a**).

**Figure 2.**
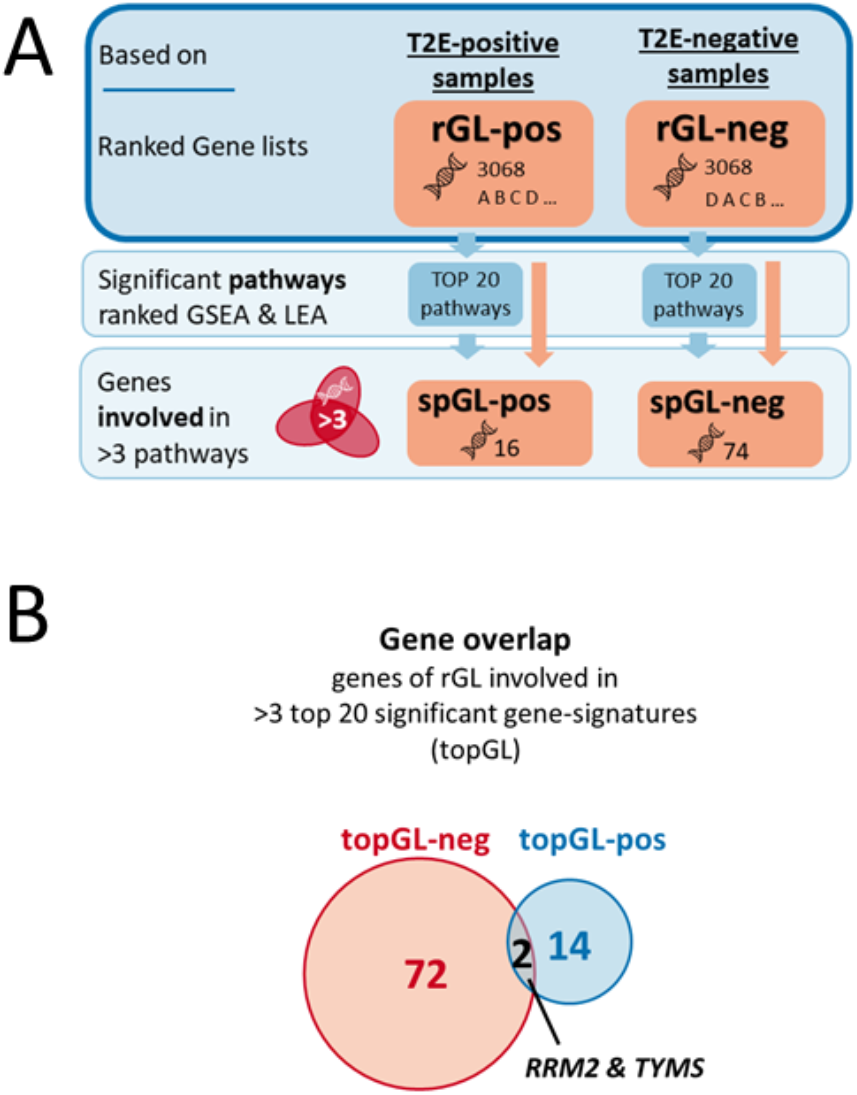
T2E-positive and -negative PCa are characterized by distinct metastasis-associated genes. **A)** Pipeline overview depicting the identification of overrepresented genes in top metastasis-associated gene-signatures in T2E-positive and -negative cases. **B)** Venn diagram showing the overlap of overrepresented genes in top metastasis-associated gene-signatures in T2E-positive and -negative cases

### Identification of genes significantly associated with metastasis

For all genes in topGL-pos and -neg the significance of differential expression in PCa patients with/without metastasis at diagnosis was determined by Mann-Whitney-U test [21]. All genes were separately tested in both PCa cohorts (TCGA-PRAD and GSE46691). *P* values were not adjusted for multiple comparison (significance for *P*<0.05). Only genes being significantly associated with metastasis in both cohorts were considered for further analyses.

### First validation cohort

For validation of survival analyses in the TCGA-PRAD-cohort, we used another GEO dataset (GSE16560) comprising 272 Swedish PCa cases with microarray expression data (6,100 genes) and corresponding clinical information including cancer-specific death and T2E-status [22].

### Survival analysis

Survival analyses were carried out in the TCGA-PRAD-cohort and the Swedish validation cohort (GSE16560) for all genes of topGL-pos and topGL-neg using the Kaplan-Meier method and the survival package of R [21,23]. For calculation of event-free survival (EFS; event = death, appearance of a new tumor, metastases, and/or relapse), both cohorts were stratified according to their intratumoral gene expression into quartiles, and *P* values were calculated using a Mantel-Haenszel test by comparing the patient groups with the most extreme gene expressions (highest versus lowest).

### Tissue microarrays (TMAs) and immunohistochemistry (IHC)

A well-characterized prostatectomy cohort comprising 133 patients with known T2E-status (**Supplementary Table 1**) diagnosed with PCa at the Institute of Pathology of the University Hospital of Bonn (Germany) was used as a second validation cohort [24]. TMAs were constructed from formalin fixed, paraffin embedded archived tissue with up to 5 cores (diameter: 1 mm) of non-necrotic tumor tissue for each patient. For IHC, RRM2 antigens were treated with ProTags IV Antigen-Enhancer (Quartett, Berlin, Germany). RRM2 was detected with a specific rabbit-anti-human RRM2 antibody (1:500, 60 min incubation time; HPA056994, Atlas Antibodies; https://www.proteinatlas.org/ENSG00000171848-RRM2/tissue), followed by an anti-rabbit IgG antibody (MP-7401 ImmPress Reagent Kit) and DAB+ chromogen (Agilent Technologies). Slides were counterstained with hematoxylin Gill’s Formula (H-3401, Vector). RRM2 immunoreactivity was quantified by an experienced data-blinded uropathologist as percentage of positive tumor cells (cytoplasmatic expression). The survMisc package for R was used for optimal cut-off selection and Kaplan-Meier survival analyses [21].

## RESULTS

### T2E-positive and -negative PCa are characterized by distinct metastasis-associated gene-signatures

T2E-positive and -negative PCa constitute distinct molecular subtypes [8–10]. To decipher molecular differences associated with metastasis in either subtype, we analyzed transcriptome profiles with matched clinicopathological data of two public cohorts (TCGA-PRAD and GSE46691). Multiple filtering steps regarding variance and regulation, and determination of the samples’ T2E-fusion status led to a unity of 3,068 variably expressed genes (see Methods). Depending on the T2E-status, we created from this set of genes two gene lists ranked by their expression fold change between patients with/without metastasis (rGL-pos and rGL-neg) (**Figure 1a**). Metastasis was chosen as a surrogate for PCa aggressiveness, because information on metastasis was available for both cohorts and usually indicates aggressiveness in PCa [4]. GSEA on rGL-pos and -neg showed no overlap between the top 20 significant metastasis-associated gene-signatures in T2E-positive and -negative cases (**Figure 1b, Supplementary Table 2**).

From those top 20 gene-signatures, we extracted genes involved in >3 of them by leading-edge analysis to create two new ‘top gene-signature’ gene lists (topGL-pos and -neg, **Figure 2a**). Accordingly, topGL-pos contained 16 genes of rGL-pos, overrepresented in significant gene-signatures of T2E-positive cases (**Supplementary Table 3**), whereas topGL-neg contained 74 genes overrepresented in significant gene-signatures of T2E-negative (rGL-neg) cases (**Supplementary Table 4**). Only two genes (*RRM2* and *TYMS*) were shared among T2E-positive and -negative cases, but involved in different gene-signatures (**Figure 2b**). Collectively, these results indicated that T2E-positive and -negative PCa are characterized by distinct metastasis-associated gene-signatures.

### Different genes are associated with metastasis in T2E-positive and -negative PCa

Next, we separately tested whether all genes of our top gene-signatures gene lists, topGL-pos and -neg (**Figure 2**), were significantly differentially expressed depending on the presence of metastasis in the TCGA-PRAD- and GSE46691-cohorts. In T2E-postive cases (topGL-pos), three genes (*GMNN, TROAP* and *WEE1*) out of 16 were significantly higher expressed (*P*<0.05) in PCa samples with metastasis. In T2E-negative cases (topGL-neg) 29 of 74 genes were significantly (*P*<0.05) higher expressed in PCa samples with metastasis. Again, we found no overlap of these significantly differentially expressed and metastasis-associated genes between T2E-positive and -negative cases (**Supplementary Tables 3** and **4**). These results further suggested that – depending on the T2E-status – distinct genes are linked to metastasis in PCa.

### Identification of subtype-specific prognostic biomarkers

To test whether the identified metastasis-associated genes were correlated with EFS, we performed Kaplan-Meier analyses in two independent cohorts. The first comprised PCa samples from TCGA-PRAD, the second was derived from another independent microarray-based study (GSE16560, first validation cohort) [22]. We only accepted genes as being associated with EFS if they were significantly (*P*<0.05) and concordantly associated with EFS in both cohorts. While none of the genes identified in screening of T2E-positive cases (topGL-pos) were consistently associated with EFS in both cohorts, seven genes were consistently associated with EFS in T2E-negative cases (*APOE, ASPN, BGN, COL1A1, LY96, RRM2* and *TYMS*). For all seven genes, patients of both cohorts had a higher risk for shorter EFS when their tumors expressed high levels of the respective genes (**Figure 3**). Interestingly, the same biomarkers showed no concordant association with EFS in T2E-positive cases. As displayed in **Table 1**, only five genes (*ASPN, BGN, COL1A1, RRM2* and *TYMS*) were associated with metastasis and EFS in all three cohorts tested, indicating that these genes could be employed for outcome prediction exclusively in T2E-negative PCa.

**Figure 3.**
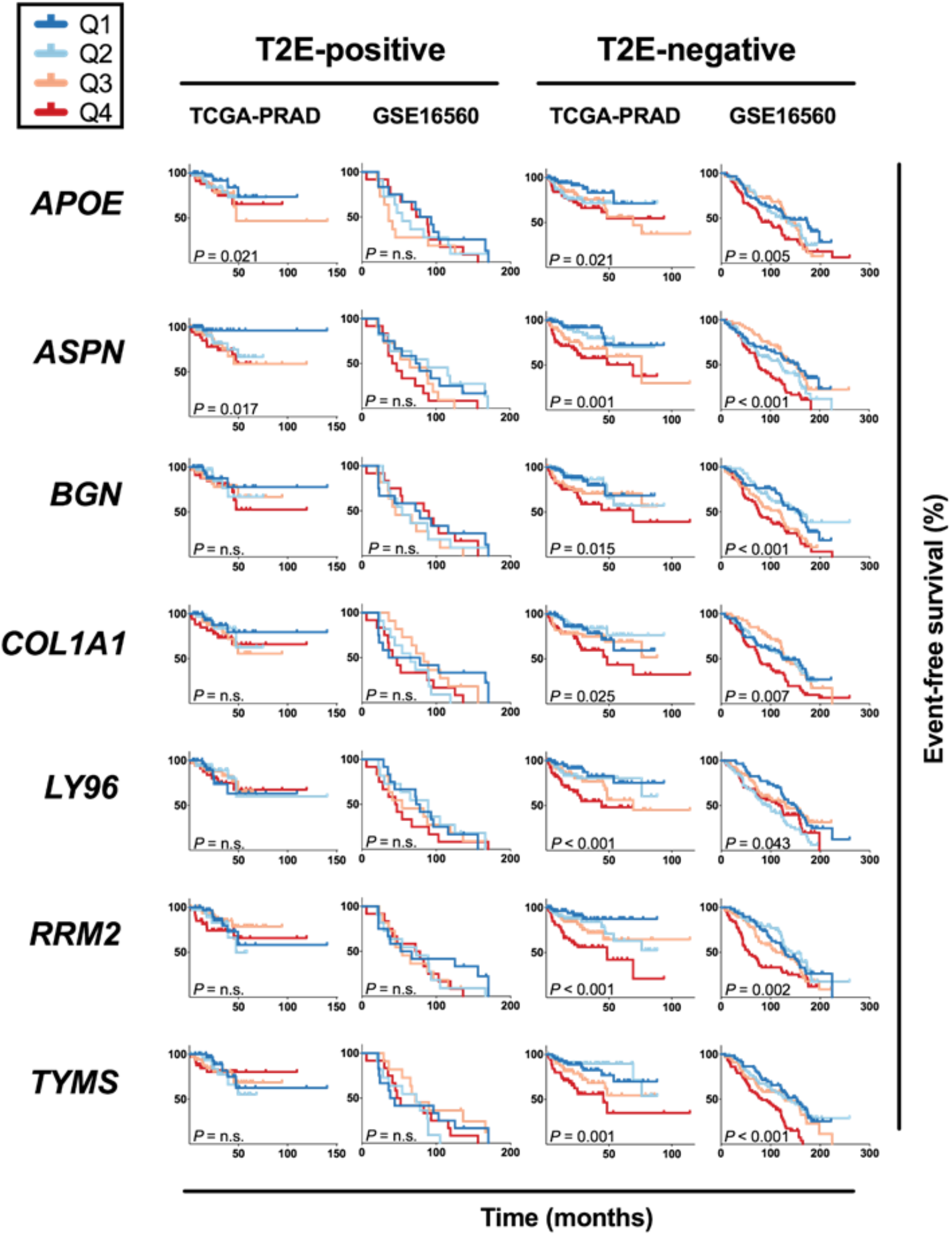
The prognostic value of identified biomarkers depends on the T2E-status. Kaplan-Meier survival plots derived from either T2E-positive or -negative samples from TCGA-PRAD- and GSE16560-cohorts for significantly event-free survival (EFS)-associated genes (*APOE, ASPN, BGN, COL1A1, LY96, RRM2* and *TYMS*) of topGL-neg. Patients were stratified by their quartile intratumoral gene expression levels of the given gene. *P* values were calculated between the lowest (Q1) and highest (Q4) gene expression quartiles using a Mantel-Haenszel test.

**Table 1.**
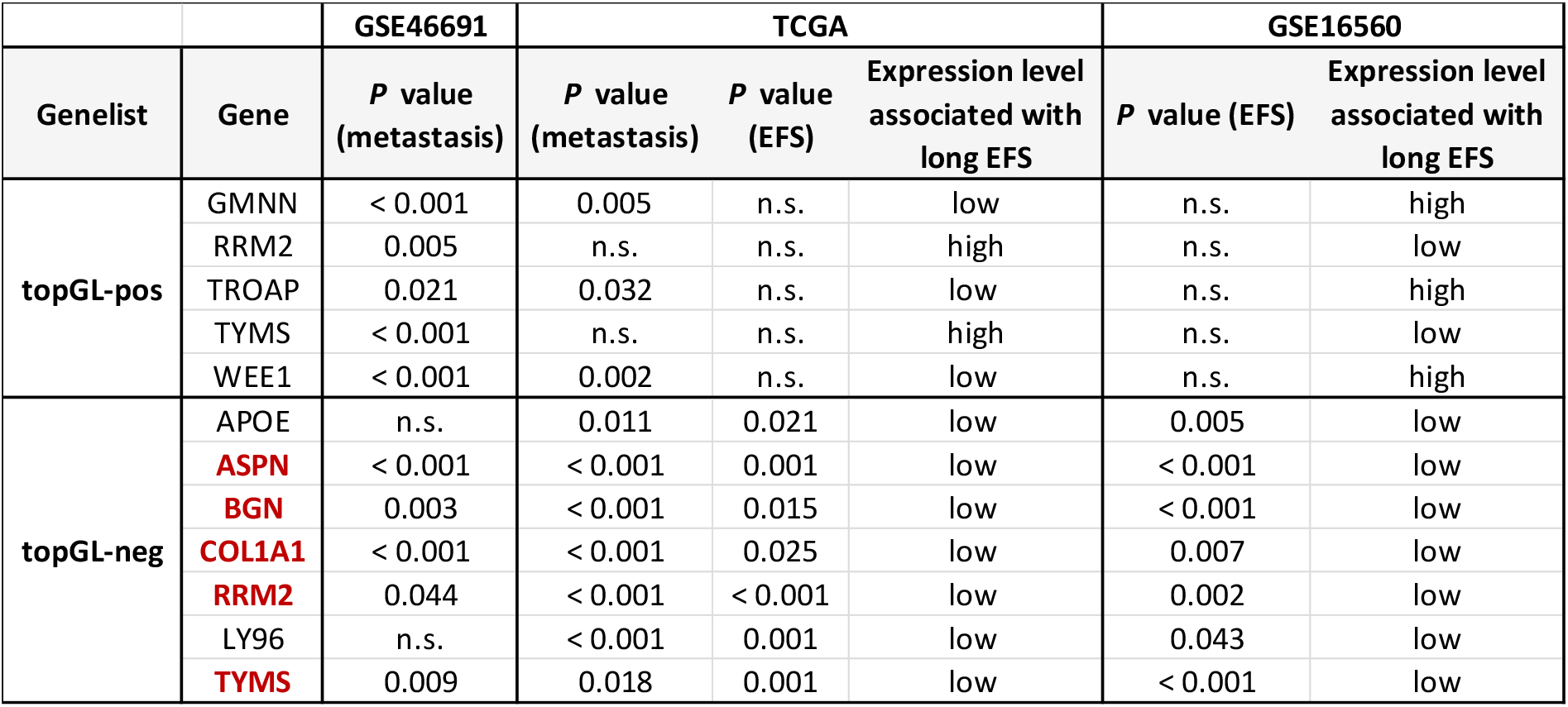
Result summary of genes in topGL-neg and topGL-pos that passed ≥1 of our tests (association test and survival analysis) for all cohorts, as well as those two genes (*RRM2, TYMS*) which were included in both gene lists (topGL-pos and -neg) (significant genes extracted from **Supplementary Tables 3 and 4**). Genes significant in all tests are highlighted in red color.

In synopsis, our results implied that the prognostic value of biomarkers critically depends on the T2E-fusion status.

### Validation of RRM2 as prognostic biomarker for T2E-negative cases by IHC

To confirm the T2E-depedent prognostic value of PCa biomarkers, we stained a TMA containing 133 PCa cases by IHC for RRM2 as an example, as for RRM2 a specific antibody was available. We separately analyzed the biochemical recurrence (BCR)-free survival of T2E-positive and -negative cases stratifying patients by their percentage of RRM2-positive tumor cells (cut-off ≥3%). We found that patients with T2E-negative PCa exhibiting a high percentage of RRM2-positive tumor cells had significantly (*P*=0.005) worse BCR-free survival than those with low RRM2-postitivity (**Figure 4**). In contrast, no association of RRM2-positivity with BCR-free survival was found in T2E-positive cases.

**Figure 4.**
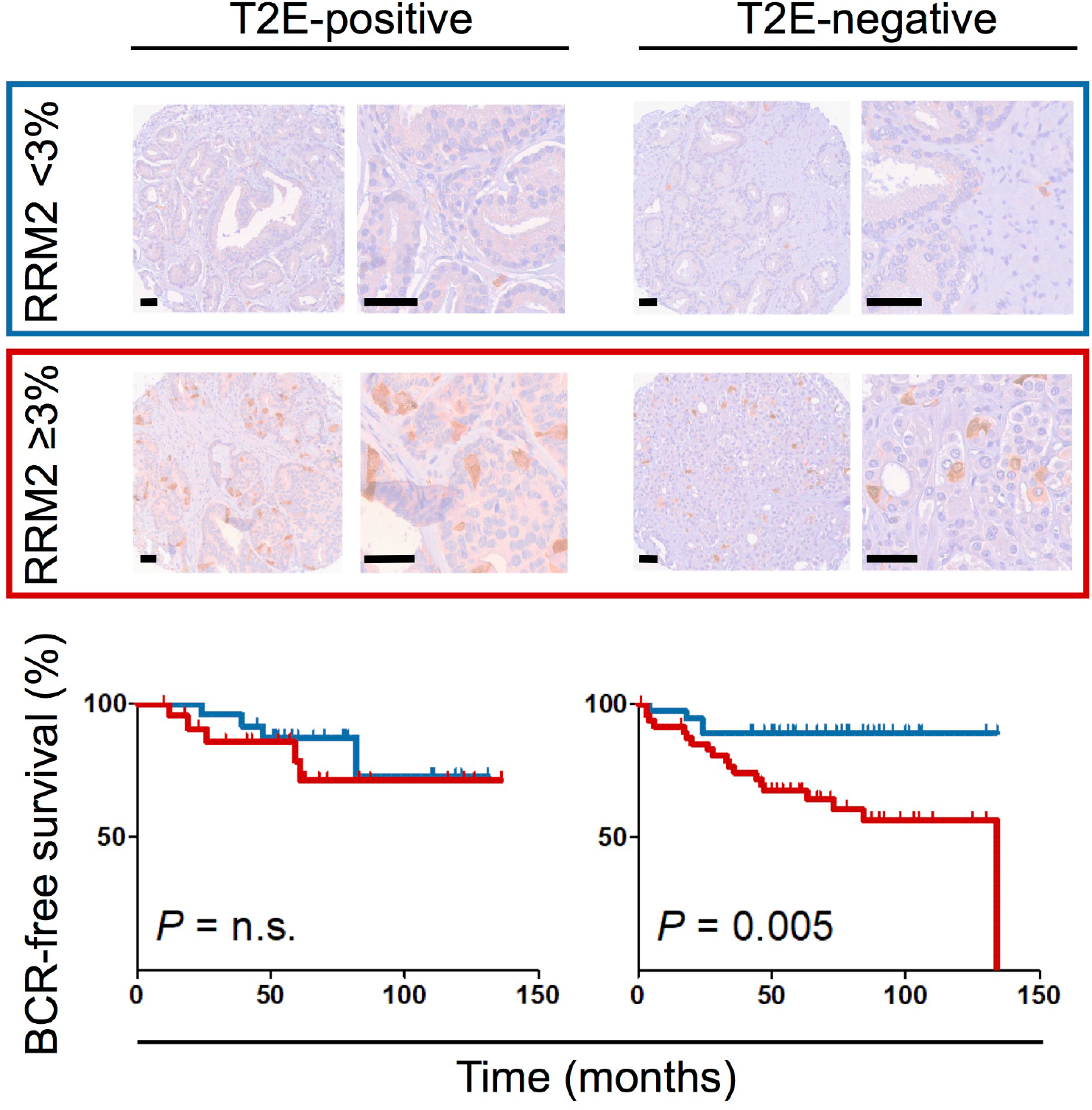
Validation of RRM2 as prognostic biomarker for T2E-negative cases by IHC. Top: Representative micrographs of T2E-positive and -negative PCa stained for RRM2 by IHC. Scale bars = 50μM for 10× and 40× magnification, respectively. Bottom: Kaplan-Meier analysis of biochemical relapse (BCR)-free survival of T2E-positive and -negative cases stratified by their median percentage of RRM2-positive tumor cells (cut-off ≥3%). Mantel-Haenszel test.

These results provided further evidence that the prognostic value of biomarkers in PCa depends on the T2E-status, and suggested that ‘pooled’ analyses ignoring the T2E-status may obscure outcome prediction.

## DISCUSSION

Prior studies showed that T2E-positive PCa are associated with specific germline susceptibility variants [8] and epigenetic profiles [9] providing evidence that T2E-positive and –negative PCa constitute distinct molecular and perhaps clinical subtypes [10]. We hypothesized that differentially expressed genes involved in distinct gene-signatures may be associated with tumor progression in T2E-positive and -negative PCa, and that prognostic biomarkers may be only relevant for a given molecular subtype.

To explore such molecular differences, we analyzed PCa transcriptomes and matched clinical data of two large cohorts (TCGA-PRAD and GSE46691). Applying several filtering steps and enrichment analyses, we identified the top 20 metastasis-associated gene-signatures for T2E-positive and -negative cases. Strikingly, these gene-signatures showed no overlap, emphasizing that T2E-positive and -negative PCa are distinct molecular subtypes that take different routes on disease progression [8–10]. From these subtype-specific gene-signatures, we extracted overrepresented genes (topGL-pos and -neg) of which five (*ASPN, BGN, COL1A1, RRM2, TYMS*) proved to be of high value exclusively for subtype-specific risk-prediction in T2E-negative PCa. These results imply that biomarkers for risk-prediction in PCa should be employed in the corresponding PCa-subtype to maximize their prognostic power.

For example, Asporin (*ASPN*) and Biglycan (*BGN*) [25] are both known to be associated with PCa progression [26] and poor prognosis [27]. Our results confirm these previous observations but highlight that they have only prognostic value for T2E-negative cases. Jacobsen *et al* additionally reported that *BGN* expression may be related to the presence of the T2E-fusion [27]. However, our study showed that in T2E-positive PCa, *BGN* is not involved in the top gene-signatures associated with metastasis, unlike in T2E-negative PCa.

The protein product of the *COL1A1* (collagen type I alpha 1), which is a major constituent of the extracellular matrix and connective tissues [25] has hitherto not been reported to be connected to outcome of PCa patients rendering COL1A1 a novel potential biomarker for T2E-negative PCa.

RRM2 (ribonuclease reductase M2) plays a role in DNA synthesis [25], and its overexpression can promote tumor progression [28]. In fact, a study not distinguishing molecular PCa-subtypes suggested that *RRM2* overexpression may be associated with PCa progression [11]. Our findings made on the mRNA and protein level are in line with these findings with the important refinement that RRM2 has strong prognostic power in T2E-negative cases, while having no prognostic value in T2E-positive cases as confirmed in four independent PCa cohorts.

Similar observations were made for *TYMS* (thymidylate synthetase), which is involved in DNA replication and repair [25] and reported to correlate with worse outcome in PCa [29]. We observed that T2E-negative patients had significantly higher risk for short EFS with high *TYMS* expression – an effect that was absent in T2E-positive cases.

In accordance with our finding that the T2E-status, which is *per se* not a strong prognostic biomarker, is crucially determining the prognostic value of other biomarkers, it has been shown that the proliferation marker Ki-67 is especially prognostic in T2E-negative cases [30,31]. However, Ki-67 (encoded by *MKI67*) was not prioritized in our screen because of discordant results in both discovery cohorts. Besides T2E-positive PCa, there are emerging additional molecular PCa-subtypes characterized by rare ETS translocations or mutations in putative driver genes such as *SPOP, FOXA1* and *IDH1* [10]. Whether these alterations also impact on biomarker prediction remains to be determined in future studies.

## CONCLUSIONS

Our findings suggest that the T2E-status should be considered when applying prognostic biomarkers to improve risk stratification of PCa patients.

## AUTHOR CONTRIBUTIONS

Julia S. Gerke and Thomas G. P. Grünewald had full access to all the data in the study and take responsibility for the integrity of the data and the accuracy of the data analysis.

*Study concept and design:* Gerke, Grünewald, Buchner

*Acquisition of data:* Gerke, Orth, Tolkach, Kristiansen, Grünewald

*Analysis and interpretation of data:* Gerke, Grünewald, Tolkach

*Drafting of the manuscript:* Gerke, Grünewald

*Critical revision of the manuscript for important intellectual content:* Buchner, Strauch, Müller-Nurasyid, Orth, Kirchner, Stief, Tolkach, Kristiansen

*Statistical analysis:* Gerke, Tolkach

*Obtaining funding:* Grünewald

*Administrative, technical, or material support:* Kirchner, Stief, Romero-Pérez, Wehweck, Stein, Musa, Knott, Hölting, Li, Sannino, Marchetto, Ohmura, Cidre-Aranaz, Kristiansen

*Supervision:* Grünewald

*Other (specify):* None

---

**Supplementary Figure 1.**
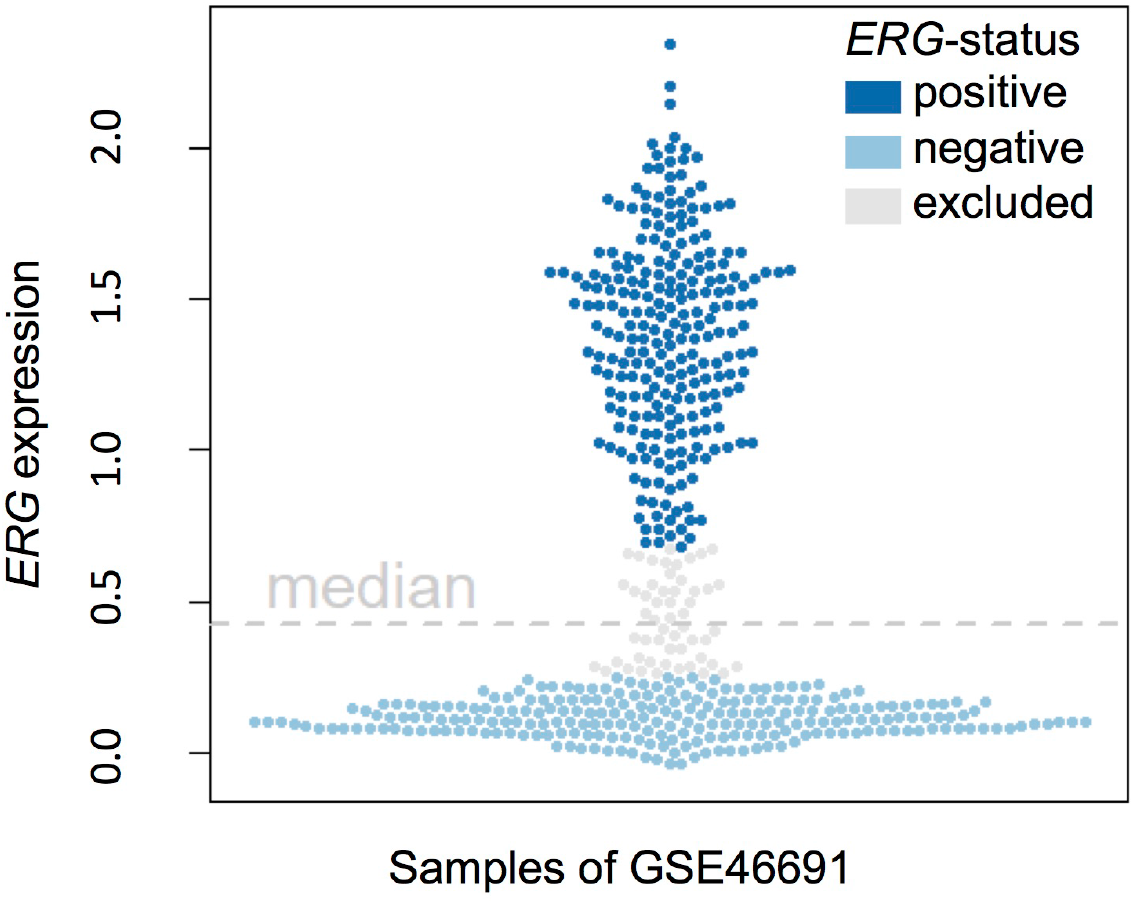
T2E-fusion status in samples of the GSE46691-cohort defined by median *ERG* expression (0.42). Samples ranging with their *ERG* expression between the 45^th^ and 55^th^ percentile (grey) were excluded.

**Supplementary Figure 2.**
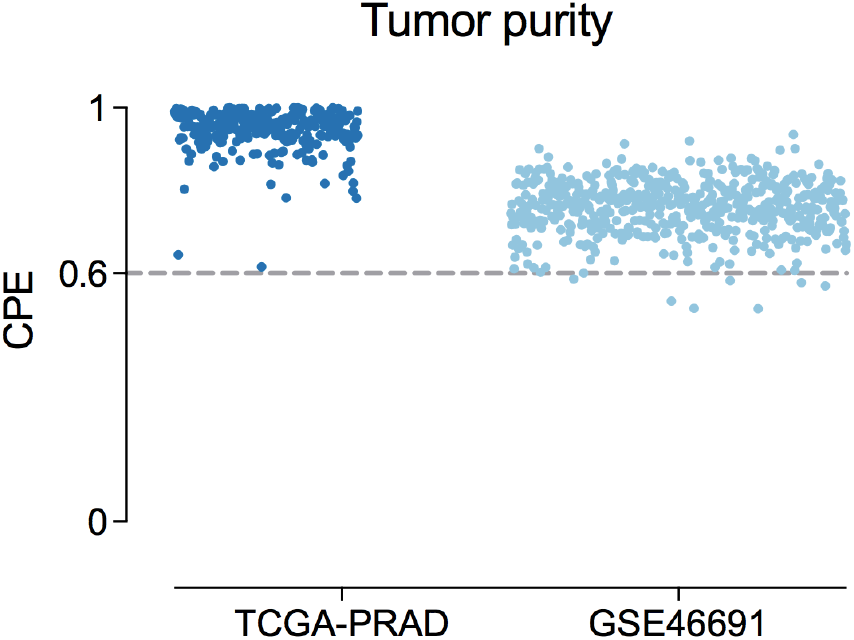
Consensus Purity Estimation (CPE) calculated with ESTIMATE algorithm of all samples of the TCGA-PRAD- and GSE46691-cohorts.

